# ABA analogue produced by *Bacillus marisflavi* modulates the physiological response of host-plant under drought stress

**DOI:** 10.1101/2020.06.12.148049

**Authors:** H.G. Gowtham, P. Duraivadivel, S. Ayusman, D. Sayani, S.L. Gholap, S.R. Niranjana, P. Hariprasad

**Affiliations:** Environmental Biotechnology Lab, Centre for Rural Development and Technology, Indian Institute of Technology Delhi, Hauz Khas, New Delhi – 110016 Delhi, India; Department of Chemistry, Indian Institute of Technology Delhi, Hauz Khas, New Delhi – 110016 Delhi, India; Department of Studies in Biotechnology, University of Mysore, Mysuru – 570006 Karnataka, India

**Keywords:** Abscisic acid, *Bacillus marisflavi*, *Brassica juncea* L., Drooping assay, Drought tolerance index, Stomatal closure

## Abstract

Present study aims to understand the molecular mechanism involved in beneficial rhizobacteria mediated alleviation of drought stress in host plant. *Bacillus marisflavi* CRDT-EB-1 isolated from the rhizosphere soil was found effective in inducing resistance against drought stress in mustard seedlings. Among the different bacterial derivatives tested, the culture filtrate was found to contain bioactive molecules. Solvent extract of bacterial culture filtrate yielded seven distinct bands/ fractions on thin layer chromatography (TLC). The fraction four (F4) with R_f_ value 0.35-0.40 was significant in reducing adverse effect of drought stress in host plants. Application of F4 resulted in delayed drooping point and higher drought tolerance index (3.34), induced stomatal closure (9.648 μ), seed germination inhibition (12%), and reduced the GA_3_ induced α-amylase activity in germinating barley seeds. On TLC, F4 turned colorless to orange color upon the spray of 2,4-dinitrophenylhydrazine reagent indicated the presence of aldehyde group. Supporting to this, the peaks between 9.8 to 10.0 ppm in ^1^H-NMR chromatogram confirmed the presence of aldehyde group. Upon LC-MS/MS analysis of crude extract of culture filtrate and F4 revealed the presence of compounds with the molecular mass 250.33 and 266.33. By analyzing these data, the identity of the bioactive compounds were predicted as xanthoxin and xanthoxic acid, which are well-known precursor of Abscisic acid (ABA) in plants. The present study concludes the capability of ABA analogue (xanthoxin like compounds) production by *B. marisflavi* CRDT-EB-1 and its involvement in inducing drought stress tolerance in the host plant.

## 1. Introduction

There is overwhelming evidence that the ancestors of modern land plants evolved in aquatic environments where they existed and diversified over millions of years. While moving from aquatic to a terrestrial environment, they come across several obstacles for which they responded through adaptive radiation, which includes several, morphological, physiological, and biochemical changes (de Vries and Archibald, 2018). Desiccation is one of the most greater challenges faced by plants for which they showed a wide range of adaptations, which include surface wax deposition and cuticles, the responsiveness of stomata to endogenous and environmental cues (Ruszala et al., 2011). Stomatal response to Abscisic acid (ABA) and CO_2_ is known to exist in lycophytes as early as 420 mYA (Rickards, 2000). During this evolutionary period, several microbes in the soil are found co-evolved with the associated plants. Moreover, the symbiotic co-evolution of plants and microbes is linked to the successful colonization of plants on land. Most of the early association is thought to facilitate the acquisition of scare and essential nutrients like phosphorus and nitrogen. With the advent of understanding the plant-microbe interaction, it was revealed that these beneficial microbes also assist the host plants to adapt under biotic and abiotic stress conditions (Ma et al., 2019; Meena et al., 2017).

The capability of beneficial rhizobacteria in inducing stress tolerance against salt and drought is well known and has been studied in the context of providing a biological means of adaptation and survival of plants under extreme abiotic stress conditions. Recent research evidences the involvement of different mechanisms in mitigating the abiotic stress in the host plant (Yang et al., 2009). The two primary mechanisms through which plant-associated beneficial bacteria known to induce tolerance against abiotic stress are by altering the level of two major stress-related hormones, ethylene and ABA. 1-aminocyclopropane-1-carboxylic acid (ACC) deaminase, an enzyme produced by plant-associated bacteria, is known to reduce the level of ethylene by degrading the ACC, an immediate precursor of ethylene (Glick, 2014). On the other hand, some of the beneficial bacteria were found inducing drought stress tolerance probably by enhancing the concentration of ABA in host plant (Cohen et al., 2009). The bacteria that have been isolated from soils, rhizosphere, endophytes, and marine samples were found capable of producing ABA in the culture defined medium (Cohen et al., 2015; Karadeniz et al., 2006; Kolb and Martín, 1985; Salomon et al., 2014; Shahzad et al., 2017). However, the induction of ABA production in host plants or the production of ABA by bacteria as a mechanism of abiotic stress management has been scarcely studied. However, the bacteria may induce resistance to abiotic stress through other mechanisms that were yet to be studied. Enhanced antioxidant enzymes such as glutathione peroxidase, superoxide dismutase etc. (Gill and Tuteja, 2010), accumulation of biocompatible solutes like proline (Ashraf and Foolad, 2007), polyphenols (Rojas-Tapias et al., 2012), tissue-specific regulation of high-affinity K^+^ transporter1 (HKT1) (Zhang et al., 2008) in host plant are some of the bacterial mediated responses involved in imparting drought and salt stress tolerance.

Reassuring to this, we found that *B. marisflavi* CRDT-EB-1 in our study was negative for ABA and ACC deaminase production. However, this isolate was found inducing drought stress tolerance in the mustard plant. Hence, the present study was designed to purify and characterize a bioactive molecule from CRDT-EB-1 responsible for the induction of tolerance against drought stress in host plants.

## 2. Materials and methods

### 2.1. Plant materials

Seeds of mustard [*Brassica juncea* L.] cultivar ‘Pusa Mustard 26 (NPJ-113)’ and Barley (*Hordeum vulgare* L.) were obtained from the Indian Agricultural Research Institute (IARI), New Delhi, India. Seeds were surface sterilized by washing with sterile distilled water, followed by rinsing with 0.5% (v/v) sodium hypochlorite (NaClO) solution for 15 min. Further, the seeds were washed with sterile distilled water for 2-3 times, blot dried, and used for all experiments. The fresh leaves of *Tradescantia fluminensis* Vell. were harvested from healthy plants naturally grown in Mahatma Gandhi Gramodaya Parisar, Indian Institute of Technology Delhi (IITD), New Delhi, India, and used for stomatal closure assay.

### 2.2. Isolation and characterization of rhizobacteria

*Bacillus marsiflavi* CRDT-EB-1 was originally isolated from the rhizospheric soil sample of mustard plants grown in the regions of Haryana (India) as explained by Hariprasad et al. (2014). The bacterial isolate was regularly sub-cultured on nutrient agar (NA) slants and maintained throughout the study. Long-term storage of bacteria was done in 40% glycerol at –86 °C.

The isolate CRDT-EB-1 was identified by following the polyphasic approaches. Gram’s staining, negative staining, and endospore staining were performed as explained by Krieg and Holt (1984). Biochemical characterization including catalase test, KOH solubility (Cappuccino and Sherman, 2013) and other 25 tests were performed using the kit (Hi-media, Bangalore) as per the manufacturer’s instruction. Further, the identity of bacterium was confirmed by amplifying and sequencing of 16S rRNA gene using universal primers and comparing it with 16S rRNA gene sequences available at National Center for Biotechnology Information (NCBI) database using the Basic Local Alignment Search Tool (BLAST) search algorithm (http://blast.ncbi.nlm.nih.gov/Blast.cgi) (Altschul et al., 1997). The nucleotide sequence was deposited into the NCBI database and obtained the accession number (MK942412). The phylogenetic tree was constructed based on 16S rRNA gene sequences by using MEGA-X software by the neighbor-joining method. The isolate CRDT-EB-1 was submitted to the National Centre for Microbial Resource (NCMR), National Centre for Cell Science (NCCS), Pune, India.

Mustard root colonizing ability was determined by following the standard procedure of Silva et al. (2003) under axenic conditions. Indole acetic acid (IAA) production was determined by growing the bacterium in Luria Bertani (LB) broth supplemented with 500 μg mL^−1^ of L–tryptophan (Patten and Glick, 2002). To analyze phosphate solubilizing ability, the bacterium was grown on the Pikovskaya medium and observed for the zone of clearance surrounding the bacterial colony (Pikovskaya, 1948). Siderophore production was determined as described by the method of Alexander and Zuberer (1991) using chrome azurol S (CAS) agar media. Nitrogen-fixing ability was determined by growing the bacteria on Burk’s N-free medium (Wilson and Knight, 1952). Hydrogen cyanide (HCN) production was determined as described by methods of Castric (1975), and Bakker and Shippers (1987) in NA amended with 4.4 g L^−1^ of glycine and 0.3 mM of FeCl_3_**·**6H_2_O and using picric acid saturated strips as indicators. Biofilm formation was qualitatively determined as described by Djordjevic et al. (2002). *In vitro* antagonistic activity against plant fungal pathogens (*Fusarium verticelloides* and *Aspergillus flavus*) was determined by following the dual culture method as described by Idris et al. (2007). Phytase production was determined by the modified methods of Fiske and Subbarow (1925), and Shimizu (1992). Method of Cattelan et al. (1999) was followed to determine the cellulase enzyme-producing ability. ACC deaminase activity of rhizobacteria was determined by the method of Penrose and Glick (2003) with minor modifications using Dworkin and Foster (DF) minimal salt broth containing 3 mM ACC as the sole N source. Antibiotic resistance was determined using octa-disc (Hi-media) with different combinations and concentrations of antibiotics (Karmakar et al., 2019). The optimum and extreme conditions (temperature, pH, salt and PEG 6000 concentration) at which the bacteria can withstand and grow was determined following standard procedures (Tiwari et al., 2018).

To determine the ABA producing ability, a loopful of 24 h old bacterial culture grown on NA was inoculated to 250 mL Erlenmeyer flask containing 100 mL of NFb chemically-defined medium (Salomon et al., 2014). The flasks were incubated at 35 ± 1 °C in an incubator shaker at 150 rpm for 2 days. After the incubation period, the bacterial growth was observed spectrophotometrically by measuring OD at 610 nm. The cultural broth was centrifuged at 8,000 rpm for 15 min at 4 °C. The supernatant was acidified to pH 2.5 with 1 N HCl and partitioned two to three times with hexane to remove lipids and other non-polar molecules. The aqueous layer was partitioned with an equal volume of ethyl acetate three times. Ethyl acetate fractions were further partitioned against the equal volume of sterile distilled water two times to remove the excess salts. The organic phase was concentrated using Rotavapor^®^ R–300 at 40 °C, and the concentrated extract was suspended in a minimal quantity of ethyl acetate. As controls, 100 mL of NFb chemically defined medium with/ without ABA (10 μg) amendment and without bacterial inoculation were maintained. Metabolites from culture filtrate and cell lysate were extracted as explained somewhere in this manuscript. The presence of ABA was determined by following the TLC, HPLC, and LC-MS methods (Antoszewski and Rudnicki, 1969; Balcke et al., 2012; Cargile et al., 1979) with minor modifications.

### 2.3. Mustard seedling drooping bioassay

*Bacillus marisflavi* CRDT-EB-1 was grown in 250 mL flask containing 100 mL of nutrient broth (NB) in a shaker for 24 h at 35 ± 1 °C. The culture was centrifuged at 8,000 rpm for 10 min at 28 ± 2 °C, and the bacterial cell pellet was suspended in sterilized distilled water. The bacterial suspension was spectrophotometrically adjusted OD_610 nm_ = 0.4-0.5. Ten g of surface-sterilized mustard seeds were bacterized for 12 h with 25 mL of bacterial suspension amended with 100 mg of carboxymethyl cellulose (CMC). Seeds soaked in sterile distilled water amended with CMC served as a control.

The experiment was conducted in a Petri dish lined with three layers of blotter discs by following the modified top of the paper (TP) method (ISTA, 2005). The bacterized and control seeds were placed equidistantly on Petri dishes and incubated in a growth chamber with controlled conditions [temperature ranging from 20 to 28 °C, around 80% of relative humidity (RH) and photoperiod of 16: 8 h (light: dark or day/ night) cycle was achieved with the fluorescent lamps (800 μmol m^−2^ s^−1^)]. Water was supplied at regular intervals equally to both control and treatment plants to maintain constant water content. The setup was incubated for 4 days in a growth chamber with the conditions as mentioned above. Before inducing the drought stress, all the plates were saturated with water. The drought stress was induced by withholding the water continuously till all the seedlings show the symptoms of drooping (a typical inverted U shaped stem) (Drooping Phase I). Watering (1 mL per Petri dish) was done, and the seedlings were allowed to recover (Recovery Phase). Further, the seedlings were allowed to droop until all the seedlings show typical drooping symptoms (Drooping Phase II). During the process of drooping, the different time points were noted, and the drooping rate, recovery rate and drought tolerance index (DTI) were calculated.

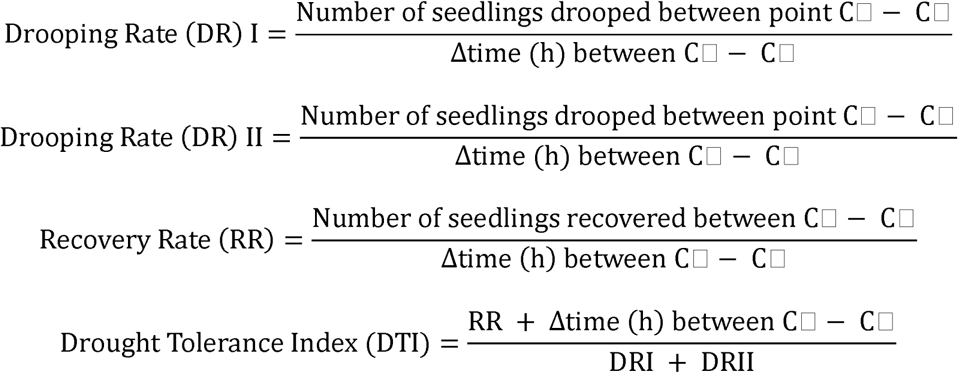

The time points selected for control experiment were explained in Fig. 1B. A similar method was followed for all different treatments. Each treatment contained three sets of four plates each, and the experiment was repeated thrice.

**Fig. 1.**
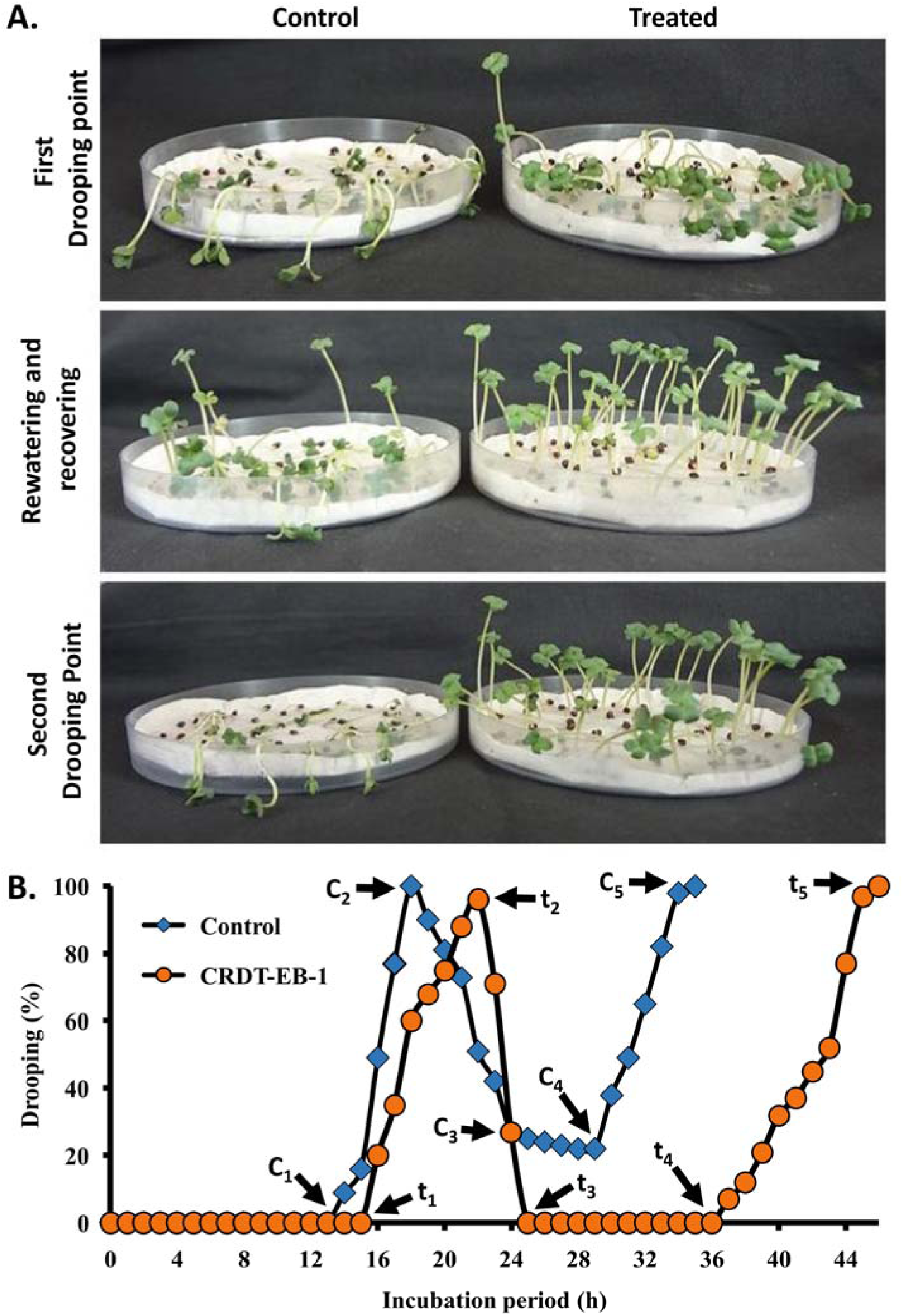
Drooping assay. (A) Effect of seed bacterization with CRDT-EB-1 on drought stress tolerance in mustard seedlings and (B) Graph representing different reading points and percentage of seedlings drooped/ recovered in control and CRDT-EB-1 treated mustard seedlings.

### 2.4. Screening of different cellular components of Bacillus marisflavi CRDT-EB-1 for their ability to induce drought stress tolerance in mustard plants

The selected bacterium was mass cultured in defined media [composed of 12.8 g L^−1^ Na_2_HPO_4_·7H_2_O; 0.3 g L^−1^ KH_2_PO_4_; 0.1 g L^−1^ NH_4_Cl as the source of N; 5 g L^−1^ beef extract; 5 g L^−1^ glucose; 2.5 g L^−1^ peptone] amended with different combinations of PEG 6000 and NaCl [2.5% PEG 6000 and 2.5% NaCl; 5% PEG 6000 and 5% NaCl; 7.5% PEG 6000 and 7.5% NaCl; 10% PEG 6000; 8% NaCl; and 10% PEG 6000 and 8% NaCl] and incubated at 35 ± 1 °C in an incubator shaker at 150 rpm for 2 days. The culture filtrate and cell pellet were separated by centrifuging the samples at 6,000 rpm for 8 min at 28 ± 2 °C. The culture supernatant was divided into two equal halves. One-half was subjected to acidification to pH 2.5 by adding 0.1 N HCl, and the crude metabolites were extracted by adding an equal amount of ethyl acetate thrice. The total protein present in another half of supernatant was precipitated by using ammonium sulfate (80%) and dialysis using membrane tubing 2 kDa molecular weight cut-off (MWCO) (Sigma-Aldrich, India).

The cell pellet collected was washed three times with phosphate buffer saline (PBS). The bacterial cells were disrupted by sonicating the suspension with 0.5 s pulses for 2 min, with an 1 min interval during which the probe was removed from the sample (Sonic Vibracell, Sonic and Materials Inc., USA). The lysed cell pellet was washed three times with PBS and used further. Cell lysate was processed similar to culture filtrate in order to obtain metabolite and protein fraction. All the cellular components and culture filtrate were used to prepare a stock solution of 1 mg mL^−1^ in distilled water. Seed treatment and drooping assay were performed as explained earlier.

### 2.5. Bioassays with rhizobacterial metabolic fractions

Mass cultivation of bacteria was performed by growing in 10 L defined media amended with 10% of PEG 6000 and 8% of NaCl under the above mentioned conditions. Collection of supernatant, processing and extraction of secondary metabolites were performed as explained above. The extract was separated on TLC Silica gel 60 F_254_ plates (Merck Life Science Pvt. Ltd., Germany) using the solvent system of hexane: isopropanol (90:10, v/v). The developed plate was visualized under short UV (254 nm) light and documented. A column chromatography (30 cm × 2 cm) was packed with 60-120 mesh size silica and the compounds were separated using gradient mobile phase containing hexane: isopropanol ranging from 100:0 to 80:20 (v/v). The fractions were analyzed through TLC, and fractions with the common band were pooled. These fractions were completely dried to remove the solvents and dissolved in distilled water at 1 mg mL^−1^ stock solution and used for different biological assays.

#### 2.5.1. Drought stress tolerance assay

The experiment was conducted in the Petri dishes lined with three layers of blotter discs following the Top of the paper (TP) method (ISTA, 2005). The surface-sterilized mustard seeds were placed equidistantly on the Petri dishes and incubated for 4 days in a growth chamber with controlled conditions, as mentioned above. Water was supplied equally to all treatments to maintain constant water content. Before inducing the drought stress, the seedlings were treated with the test solutions containing different concentrations of the bioactive fraction (ranging from 0 to 100 µg mL^−1^, prepared in distilled water) and ABA (100 µM). Treatment was done by flooding the Petri dishes with 1 mL of test solutions. Seedlings treated with the same quantity of distilled water served as control. The experimental setup was incubated under earlier said conditions. DR-I, DR-II, RR and DTI were determined as explained earlier. Each treatment contained three sets, each containing four Petri dishes, and the experiment was repeated thrice.

#### 2.5.2. Stomatal closure assay

Stomatal closure assay was carried out as described by Wang et al. (2001) by treating the leaves of *T. fluminensis* with different column fractions of bacterial extract and ABA. Briefly, fresh leaves of *T. fluminensis* were harvested and cut into a piece of 1 cm^2^, avoiding the mid-vein. To induce the stomatal opening, the adaxial surface of leaf pieces were placed downward in the Petri dishes (90 mm diameter) containing the buffer (composed of 5 mM MES hydrate buffer, pH 6.15, 20 mM KCl and 1 mM CaCl_2_·2H_2_O) at 25 °C for 3 h under the illumination at 200 µmol m^−2^ s^−1^. Further, these leaf pieces were transferred to fresh buffer containing different fractions of bacterial extract and ABA and then incubated for 1 h. After the incubation period, the transparent nail polish was applied immediately to the lower surface of the leaf and allowed to dry. Then, the clear transparent packing tape was used to peel off the nail polish, which was then placed onto a plain microscope slide. Images were captured by Leica microscopic model 750, which attached to a digital camera system model EC3 (Leica Microsystems, Germany). The stomatal aperture measurements were performed for each treatment using ImageJ 1.47 v software (National Institutes of Health, Bethesda, USA). The ABA (2.5 and 25 μM) and distilled water were used as positive and negative controls, respectively.

#### 2.5.3. Seed germination inhibition assay

Seed germination inhibition assay was carried out by soaking the surface-sterilized mustard seeds for 12 h with the different column fractions of bacterial extract (100 µg mL^−1^) and ABA (25 μM). Further, the seeds were treated with different concentrations of F4 ranging from 10, 100, and 1000 μg mL^−1^, and ABA (6.25, 12.5 and 25 μM) treatment was also maintained as control. The treated and control seeds were plated in the Petri plates lined with three layers of blotting sheet saturated with water. The setup was incubated at 28 ± 2 °C for 3 days, and the germination percentage was calculated. For each treatment, four plates were maintained with 50 seeds each, and the experiment was repeated thrice.

#### 2.5.4. Inhibition of gibberellic acid (GA_3_) induced α-amylase activity

Quantification of α-amylase activity was carried out by the method as described by Ho et al. (1980). Barley seeds were cut transversely to remove embryo portions, resulting in half-seeds without embryos. Half-seeds were then surface sterilized as described above. Ten barley half-seeds were placed in 2 mL of imbibing solution (20 mM sodium succinate, pH 5.0, and 20 mM CaCl_2_) with various hormones (GA_3_ and ABA) and the test compound F4 at different concentrations. Treatments and concentrations of hormones were as follows: (a) No hormone; (b) GA_3_ (1 µM); (c) ABA (20 µM); (d) F4 (100 µg mL^−1^); (e) GA_3_ (1 µM) + ABA (20 µM); and (f) GA_3_ (1 µM) + F4 (100 µg mL^−1^). Then, the seeds placed in imbibing solutions containing various hormones were incubated at 24 °C for 28 h. After incubation, the enzyme solution was harvested and tested for the α-amylase activity using a spectrophotometer. Briefly, 15 mg potato starch was added to 10 mL of imbibing solution. The solution was boiled for 1 min and then centrifuged at 10,000 rpm for 15 min at 4 °C. Next, 50 µL of supernatant was mixed with 100 µL of enzyme solution and incubated for 180 min, and then reacted with 100 µL of I_2_/KI solution (23.6 mM I_2_ and 0.36 M KI dissolved in 0.1 N HCl). The absorbance was read at 620 nm using a spectrophotometer. The α-amylase activity was calculated using the following formula:

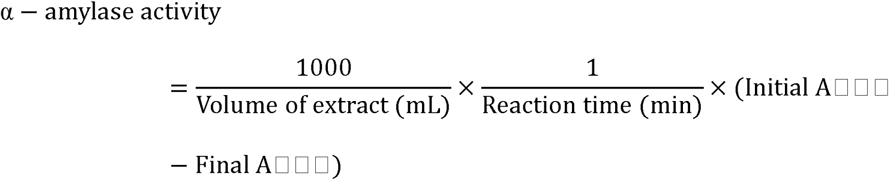

Then, the α-amylase activity was compared with the control samples to calculate induction and suppression values. The activity was expressed as ΔA_620_ min^−1^ mL^−1^.

#### 2.5.5. Reversal of fluridon induced drought stress in mustard seedlings

Mustard seeds were plated on blotter sheets saturated with 100 µM fluridon (Sigma-Aldrich, USA) solution as explained earlier. The setup was incubated in the growth chamber (under dark conditions) at 22 °C. Similarly, a distilled water-treated control was maintained. Four days old-seedlings were removed from dark conditions and treated by flooding the plates with 1 mL of distilled water or F4 (100 µg mL^−1^) or ABA (10 µM). The plates were allowed to dry, and % drooping was calculated as explained earlier.

### 2.6. Characterization of bioactive compounds present in F4

The crude extract and purified fractions were separated on TLC Silica gel 60 F_254_ plates (Merck Life Science Pvt. Ltd., Germany) using the solvent system of hexane: isopropanol (90:10, v/v). The developed plate was visualized under short UV (254 nm) light and documented. A 2,4-dinitrophenylhydrazine reagent was freshly prepared as described by Ruekberg and Rossoni (2005) and sprayed on the developed plate to visualize the presence of aldehyde group as orange color band. LCMS analysis was performed using mass spectrometer (Q-Exactive Plus Biopharma-High Resolution Orbitrap) coupled to HPLC equipped with UV–Visible detector (SAIF, IIT Bombay, India). The mobile phases were 0.1% formic acid in water (A) and 90% acetonitrile in water with 0.1% formic acid (B). The LC conditions were 5% B during 0–3 min, a linear increase from 5% to 20% B during 3–25 min, from 20% to 40% B during 25–40 min and from 40% to 50% B during 40–55 min, finally from 50% to 95% B during 55–63 min followed by 15 min of maintenance. Injection volume 8 μL and flow rate 0.4 mL min^−1^. ESI parameters: both positive and negative modes. The scan was collected in the Orbitrap at a resolution of 30,000 in an *m/z* range of 50–1500. ^1^H-NMR analysis of F4 was carried out on a Bruker Avance 500 AV spectrometer operating at 500 MHz (IIT Delhi, India). The sample was dissolved in CDCl_3_ and measured with a spectral width of ∼ 10,000 Hz, acquisition time of 3.27 s and a relaxation delay of 1.0 s was applied.

### 2.7. Statistical analysis

All data were statistically analyzed and subjected to arcsine transformation and analysis of variance (ANOVA) using SPSS, version 23 (SPSS Inc., Chicago, IL). The significant differences between the treatment means were compared using the Highest Significant Difference (HSD) as obtained by Tukey’s test at *P* ≤ 0.05 level.

## 3. Results

### 3.1. Characterization of Bacillus marisflavi

Isolate CRDT-EB-1 was morphologically dark to pale yellow in color. The colonies on NA were round with even margin doom shaped and shining (Supplementary Fig. S1). The isolate was Gram +ve or Gram-variable, rod-shaped and endospore-forming. Biochemical characters recorded by the isolate were summarized in Supplementary Table S1. Further, the identity of the bacterial isolate was confirmed by comparing its 16S rRNA gene sequence with the existing sequence at the NCBI database. The sequence was submitted to the NCBI database, and the accession number was obtained (MK942412). By analyzing all the results obtained, the bacterial isolate was identified as *Bacillus marisflavi*. The isolate CRDT-EB-1 was deposited to the NCMR-NCCS, Pune, India with the accession number MCC 4286.

The bacterial isolate was found growing in the temperature range of 10-45 °C (optimum: 30-37 °C); pH 3-11 (optimum: 5-9); NaCl 0-15% (optimum: 0-8%) and PEG 6000 0-25% (optimum: 0-10%). The beneficial traits of *B. marisflavi* CRDT-EB-1 were summarized in Table 1. The bacterial isolate was able to colonizing the root of mustard and found positive for nitrogen-fixing ability, IAA and siderophore production. However, the isolate was negative for phosphate solubilization, HCN production, biofilm formation, antifungal activity, phytase, cellulase, ACC deaminase activity, and ABA production (Table 1, Supplementary Fig. S2 and S3). Seed germination, root length, shoot length and seedling vigor were not affected upon seed treatment with CRDT-EB-1 (Table 1).

**Table 1.** Plant growth promoting traits of *Bacillus marisflavi* CRDT-EB-1.

| Characters | Results |
| --- | --- |
| Root colonization | + |
| IAA production | + |
| Phosphate solubilization | – |
| Siderophore production | + |
| Nitrogen fixation | + |
| HCN production | – |
| Biofilm formation | – |
| Antifungal activity | – |
| Phytase production | – |
| Cellulase production | – |
| ACC deaminase activity | – |
| ABA production | – |
| Plant growth promoting activity<br>(% increase over control) |  |
| Seedling vigor | 17.9 |
| Root length | 20.0 |
| Shoot length | 12.5 |
| Germination | 2.22 |

‘+’ indicates positive and ‘–’ indicates negative.

The production of extracellular metabolites was influenced by different concentrations of PEG 6000 and NaCl amendment to the culture media (Supplementary Fig. S4). Among the various combination of media used to grow a bacterium, the culture filtrate metabolic fraction of minimal salt broth amended with 10% of PEG 6000 and 8% of NaCl was found significant in delaying drooping of mustard seedlings followed by culture filtrates of defined medium amended with 7.5% PEG 6000 and 7.5% NaCl, and 5% PEG 6000 and 5% NaCl (Data not shown).

Results of drooping assay clarified that both bacterium and culture filtrate - MF (minimal salt broth amended with 10% of PEG 6000 and 8% of NaCl) could induce drought resistance to various extent, which was evidenced by delay in DR-I and DR-II and improved RR (Fig. 1, Table 2). Also, the time between two drooping points was significantly high (26 h) in comparison to control (17 h). The DTI was significantly improved from 0.860 in control to 2.362 for bacterial treated seedlings and 2.881 for culture filtrate - MF treated seedlings. The cell lysate - MF also recorded higher DTI (1.847), and other treatments were almost equivalent to control in all parameters studied (Table 2).

**Table 2.**
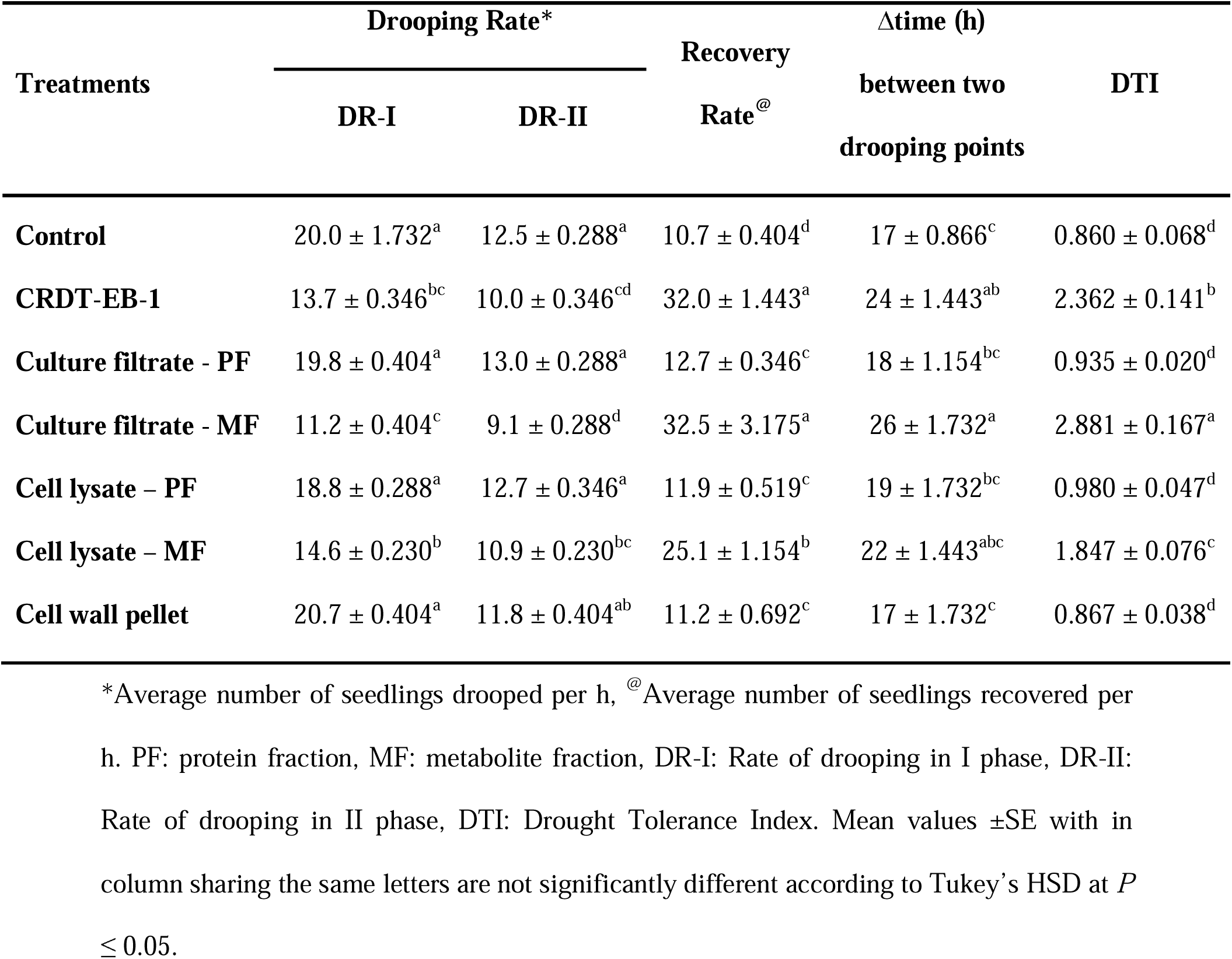
Effect of seedling treatment with CRDT-EB-1 and their different cultural fractions on drought stress tolerance ability of mustard seedlings.

| Treatments | Drooping Rate* | | Recovery<br>Rate<sup>@</sup> | Δtime (h)<br>between two<br>drooping points | DTI |
| --- | --- | --- | --- | --- | --- |
|  | DR-I | DR-II |  |  |  |
| **Control** | 20.0 ± 1.732<sup>a</sup> | 12.5 ± 0.288<sup>a</sup> | 10.7 ± 0.404<sup>d</sup> | 17 ± 0.866<sup>c</sup> | 0.860 ± 0.068<sup>d</sup> |
| **CRDT-EB-1** | 13.7 ± 0.346<sup>bc</sup> | 10.0 ± 0.346<sup>cd</sup> | 32.0 ± 1.443<sup>a</sup> | 24 ± 1.443<sup>ab</sup> | 2.362 ± 0.141<sup>b</sup> |
| **Culture filtrate - PF** | 19.8 ± 0.404<sup>a</sup> | 13.0 ± 0.288<sup>a</sup> | 12.7 ± 0.346<sup>c</sup> | 18 ± 1.154<sup>bc</sup> | 0.935 ± 0.020<sup>d</sup> |
| **Culture filtrate - MF** | 11.2 ± 0.404<sup>c</sup> | 9.1 ± 0.288<sup>d</sup> | 32.5 ± 3.175<sup>a</sup> | 26 ± 1.732<sup>a</sup> | 2.881 ± 0.167<sup>a</sup> |
| **Cell lysate – PF** | 18.8 ± 0.288<sup>a</sup> | 12.7 ± 0.346<sup>a</sup> | 11.9 ± 0.519<sup>c</sup> | 19 ± 1.732<sup>bc</sup> | 0.980 ± 0.047<sup>d</sup> |
| **Cell lysate – MF** | 14.6 ± 0.230<sup>b</sup> | 10.9 ± 0.230<sup>bc</sup> | 25.1 ± 1.154<sup>b</sup> | 22 ± 1.443<sup>abc</sup> | 1.847 ± 0.076<sup>c</sup> |
| **Cell wall pellet** | 20.7 ± 0.404<sup>a</sup> | 11.8 ± 0.404<sup>ab</sup> | 11.2 ± 0.692<sup>c</sup> | 17 ± 1.732<sup>c</sup> | 0.867 ± 0.038<sup>d</sup> |

\*Average number of seedlings drooped per h, <sup>@</sup>Average number of seedlings recovered per h. PF: protein fraction, MF: metabolite fraction, DR-I: Rate of drooping in I phase, DR-II: Rate of drooping in II phase, DTI: Drought Tolerance Index. Mean values ± SE with in column sharing the same letters are not significantly different according to Tukey's HSD at $P \leq 0.05$ .

### 3.2. Bioassay-guided purification of bioactive compound

On the TLC plate, separation of crude extract of bacterial culture filtrate yielded 7 distinct bands with the mobile phase hexane: isopropanol (90:10, v/v). All the 7 bands (fractions) were collected through the column chromatography and subjected to bioassays. The fraction 4 (F4) corresponding to the R_f_ value 0.35-0.40 and turned to orange color upon the spray of 2,4-dinitrophenylhydrazine reagent on the TLC plate recorded significant improvement in drought stress tolerance as evident by increased DTI (3.344) in comparison to control and other bands (Table 3). In dose-dependent drooping assay, different concentrations of F4 applications revealed that the maximum of DTI of 3.238 was reached at 10 μg mL^−1^ concentration beyond that the DTI was not significantly different (Fig. 3). Further, F4 showed to induce the stomatal closure in the leaves of *T. fluminensis* with least stomatal aperture measured (9.648 µ) in comparison to other fractions and control (27.116 µ) (Fig. 4). In the case of seed germination inhibition assay, the mustard seeds treated with F4 (10 µg mL^−1^) recorded 99% inhibition, which was equivalent to ABA (12.5 µM) treatment (Fig. 5). Other fractions were not significant in inhibiting seed germination. The α-amylase activity of untreated barley seeds was found to 47.88 U. Upon treating with GA_3_, the activity was increased to 62.61 U. The F4 was found to suppress the α-amylase activity (33.55 U) greater than ABA treatment (41.27 U). Similarly, the most significant decrease in α-amylase activity was observed in GA_3_ + F4 treatment (21.27 U) (Fig. 6). The application of F4 reversed the fluridon induced drought stress in mustard seedlings (Fig. 7). The drooping rate was significantly reduced by F4, which was in accordance with the effect of ABA observed in fluridon treated seedlings.

**Table 3.** Effect of seedling treatment with different column fractions of CRDT-EB-1 culture supernatant on drought stress tolerance ability of mustard seedlings.

| Treatments | Drooping Rate* | | Recovery<br>Rate<sup>@</sup> | Δtime (h)<br>between two<br>drooping<br>points | DTI |
| --- | --- | --- | --- | --- | --- |
|  | DR-I | DR-II |  |  |  |
| **Control** | 20.0 ± 0.288<sup>ab</sup> | 12.5 ± 1.212<sup>a</sup> | 10.7 ± 0.750<sup>b</sup> | 17 ± 1.443<sup>b</sup> | 0.852 ± 0.028<sup>c</sup> |
| **F1** | 19.7 ± 0.404<sup>ab</sup> | 12.8 ± 0.288<sup>a</sup> | 11.0 ± 0.866<sup>b</sup> | 18 ± 1.154<sup>b</sup> | 0.892 ± 0.036<sup>c</sup> |
| **F2** | 20.8 ± 0.692<sup>a</sup> | 13.1 ± 0.404<sup>a</sup> | 10.1 ± 0.461<sup>b</sup> | 17 ± 1.732<sup>b</sup> | 0.800 ± 0.056<sup>c</sup> |
| **F3** | 20.1 ± 0.635<sup>ab</sup> | 12.6 ± 0.519<sup>a</sup> | 11.2 ± 0.635<sup>b</sup> | 17 ± 1.443<sup>b</sup> | 0.862 ± 0.034<sup>c</sup> |
| **F4** | 10.2 ± 0.404<sup>c</sup> | 8.4 ± 0.346<sup>c</sup> | 34.2 ± 0.750<sup>a</sup> | 28 ± 1.732<sup>a</sup> | 3.344 ± 0.195<sup>b</sup> |
| **F5** | 19.8 ± 0.461<sup>ab</sup> | 12.2 ± 0.404<sup>a</sup> | 10.9 ± 1.096<sup>b</sup> | 18 ± 1.443<sup>b</sup> | 0.903 ± 0.028<sup>c</sup> |
| **F6** | 19.6 ± 0.923<sup>ab</sup> | 12.9 ± 1.385<sup>a</sup> | 10.1 ± 0.808<sup>b</sup> | 18 ± 2.020<sup>b</sup> | 0.864 ± 0.024<sup>c</sup> |
| **F7** | 17.0 ± 1.154<sup>b</sup> | 11.0 ± 1.154<sup>ab</sup> | 14.3 ± 0.750<sup>b</sup> | 20 ± 1.443<sup>b</sup> | 1.225 ± 0.130<sup>c</sup> |
| **ABA (25 μM)** | 8.6 ± 0.519<sup>c</sup> | 7.8 ± 0.404<sup>c</sup> | 35.5 ± 2.482<sup>a</sup> | 28 ± 1.732<sup>a</sup> | 3.871 ± 0.195<sup>a</sup> |

\*Average number of seedlings drooped per h, <sup>@</sup> Average number of seedlings recovered per h. DR-I: Rate of drooping in I phase, DR-II: Rate of drooping in II phase, DTI: Drought Tolerance Index. Mean values ± SE with in column sharing the same letters are not significantly different according to Tukey's HSD at $P \leq 0.05$ .

**Fig. 2.**
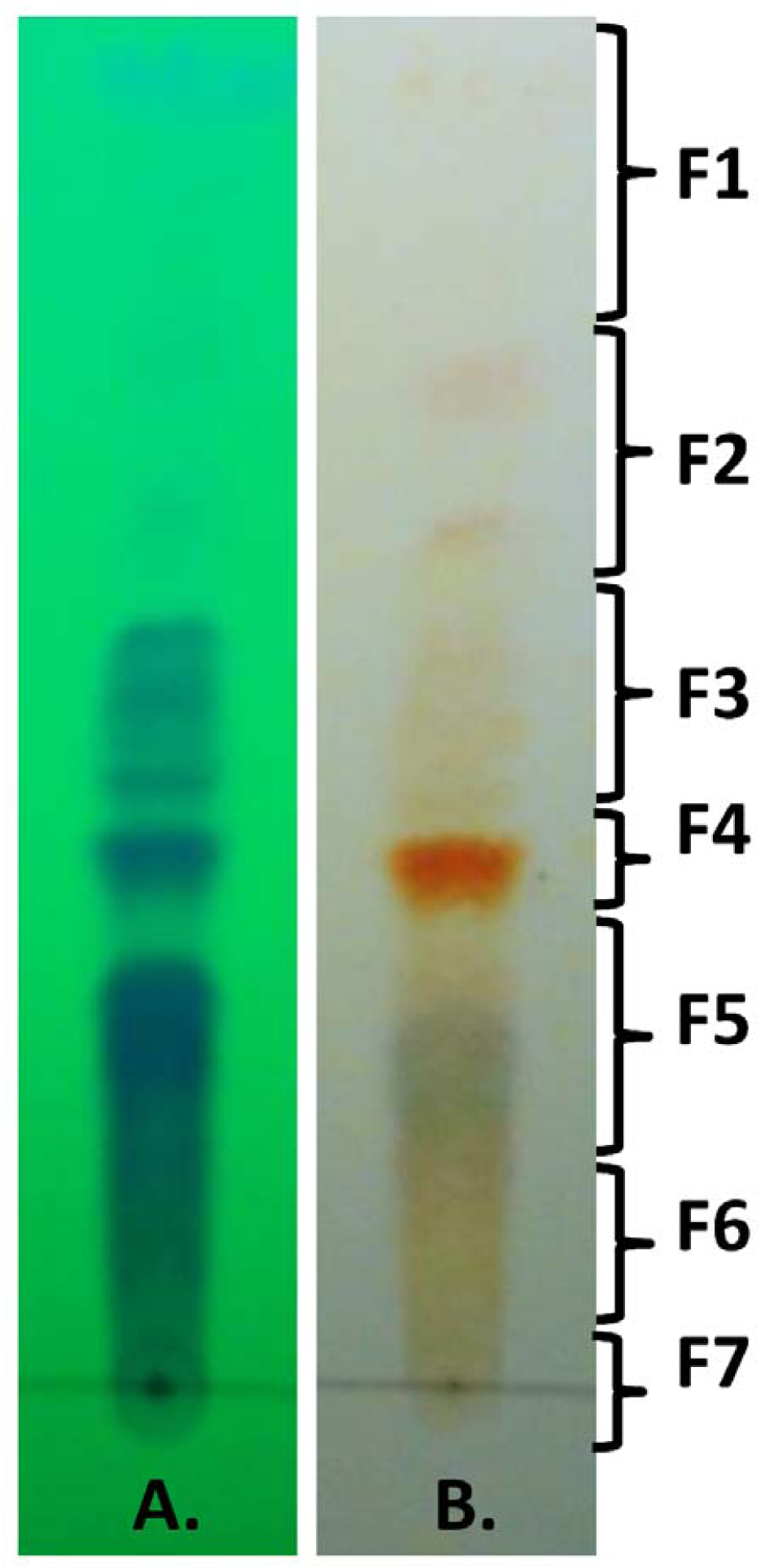
Thin layer chromatographic separation of metabolites present in crude extract of CRDT-EB-1 culture supernatant using mobile phase hexane: iso-propanol (90:10, v/v) (A) observed under short UV (254 nm) light and (B) after spraying with 2,4-dinitrophenylhydrazine reagent. F1 – F7 are the fractions used for bioassay.

**Fig. 3.**
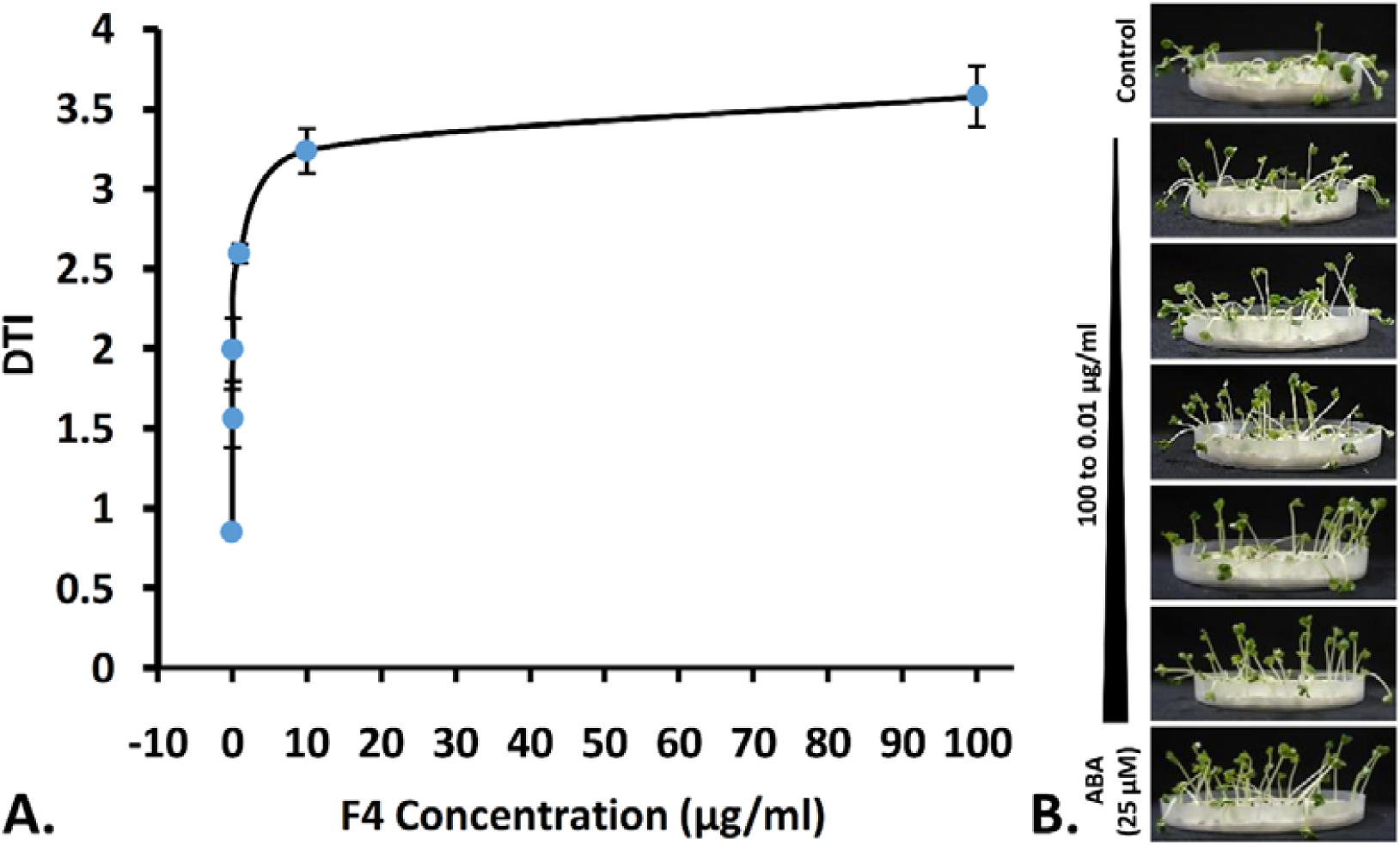
Effect of seedling treatment with different concentrations of F4 on drought tolerance index (DTI) of mustard seedlings.

**Fig. 4.**
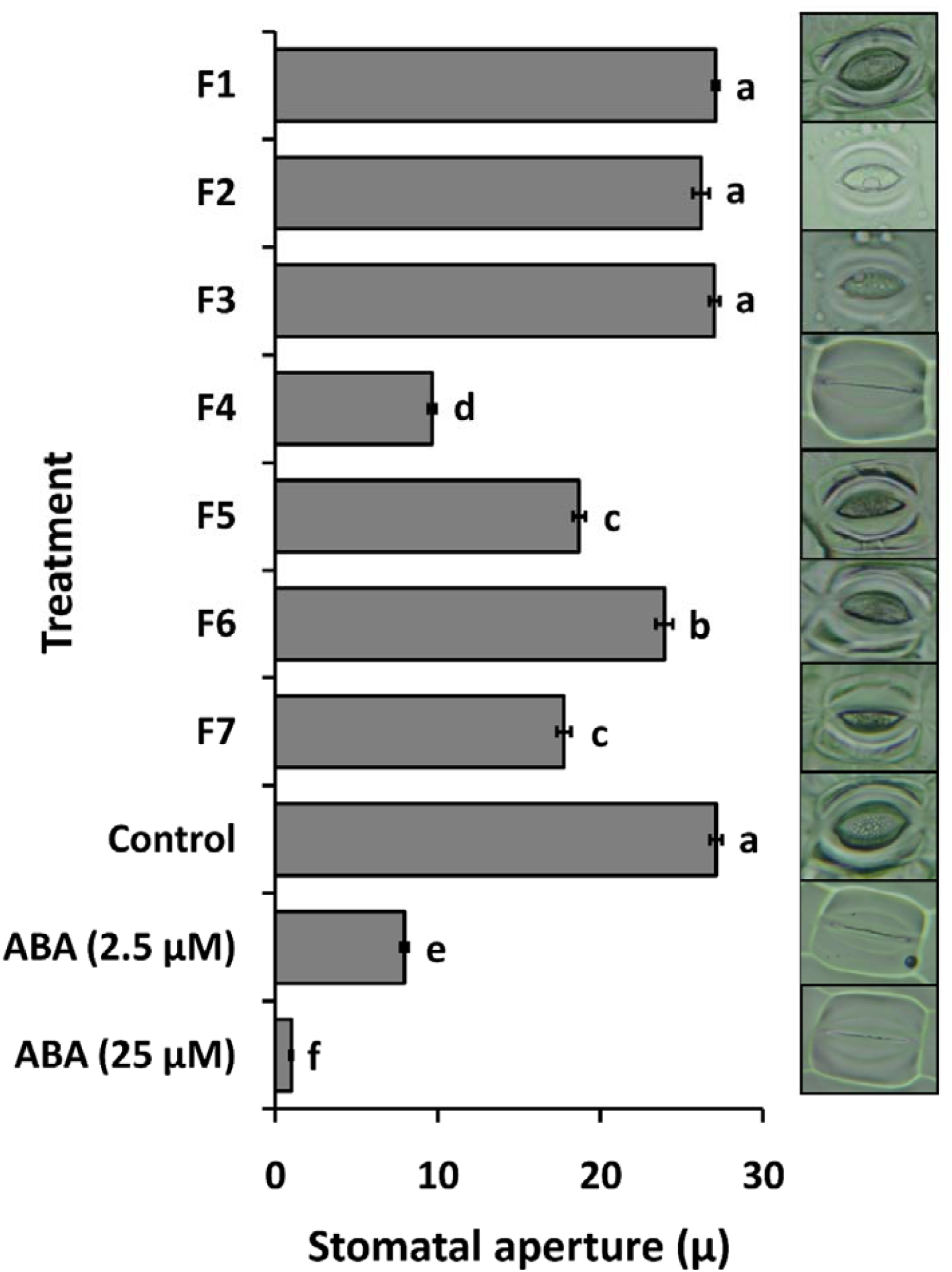
Induction of stomatal closure by different column fractions of CRDT-EB-1 culture filtrate crude extract at 100 µg mL^−1^ concentration.

**Fig. 5.**
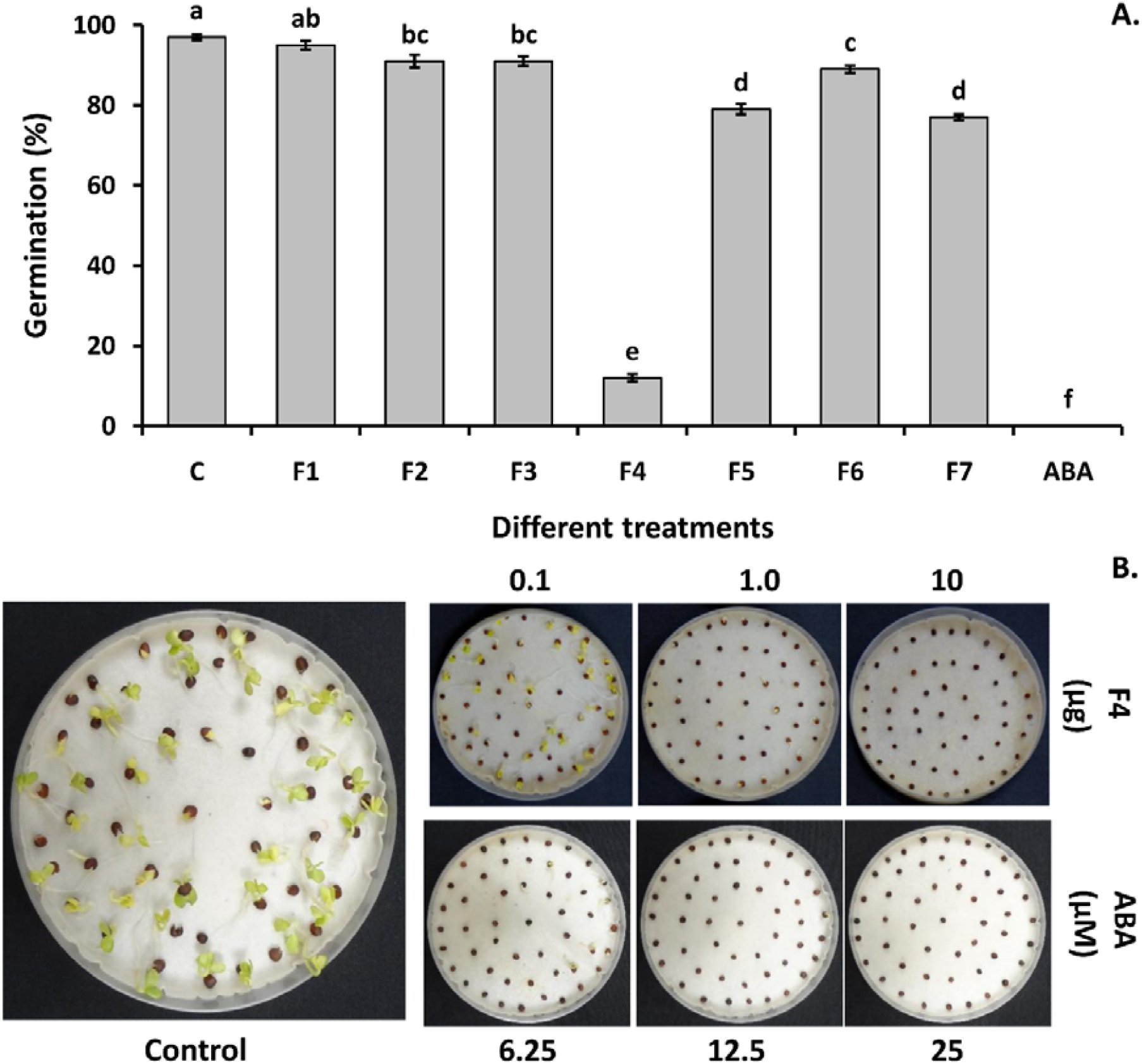
Seed germination inhibition bioassay. (A) Inhibition of mustard seed germination by different fractions of CRDT-EB-1 culture filtrate crude extract at 100 µg mL^−1^ concentration, and (B) Inhibition of mustard seed germination by different concentrations of F4 as compared with ABA.

**Fig. 6.**
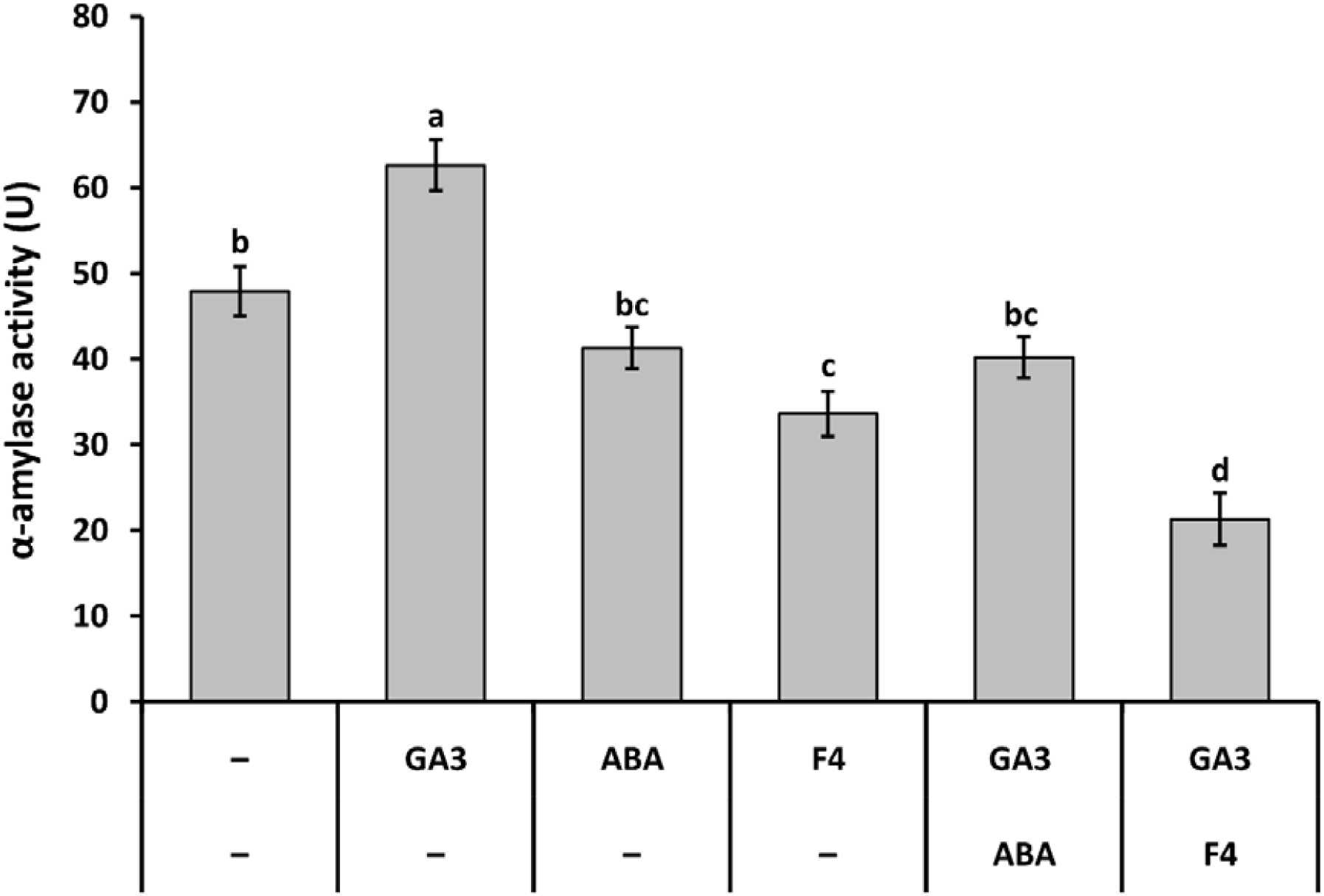
Inhibition of GA_3_ induced α-amylase activity in barley seeds by F4 (100 µg mL^−1^) and ABA (20 μM).

**Fig. 7.**
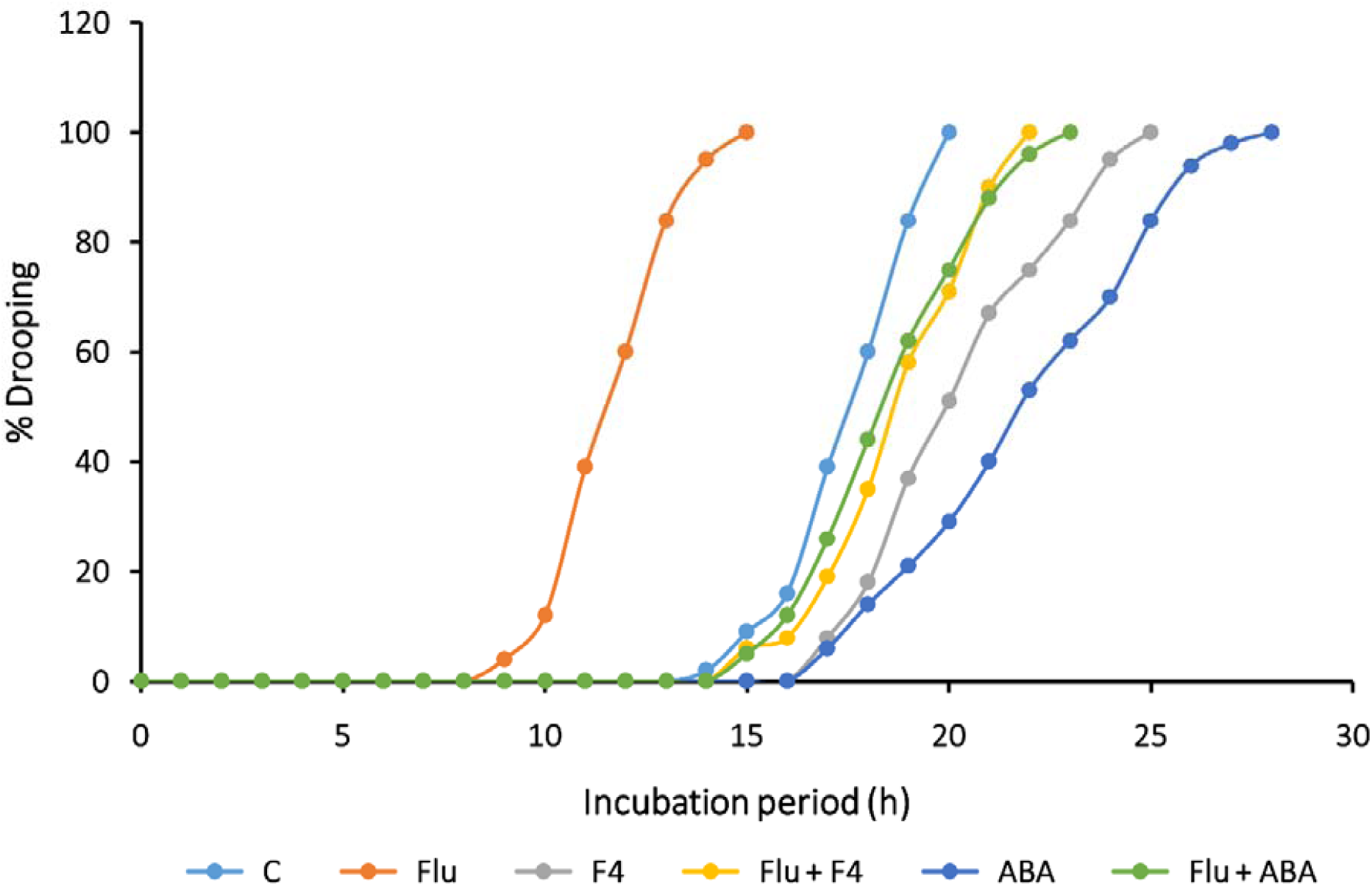
Reversal of fluridon induced accelerated drooping by F4 and ABA in mustard seedlings.

### 3.3. Identification of bioactive compounds present in F4

Upon separating the crude metabolites of culture filtrate on TLC followed by spraying with 2,4-dinitrophenylhydrazine reagent resulted in color change from invisible to orange, indicated the presence of aldehyde group in F4. Further, LC-MS analysis revealed the presence of multiple compounds in F4. Among them, two compounds with molecular mass of 250.12 and 266.21 Da were reported which were suspected as xanthoxin and xanthoxic acid (Fig. 8A). Supporting to this, the presence of peaks between 9.8 to 10.0 ppm (Fig. 8B) in ^1^H-NMR indicated the presence of aldehyde group, a structural component of xanthoxin.

**Fig. 8.**
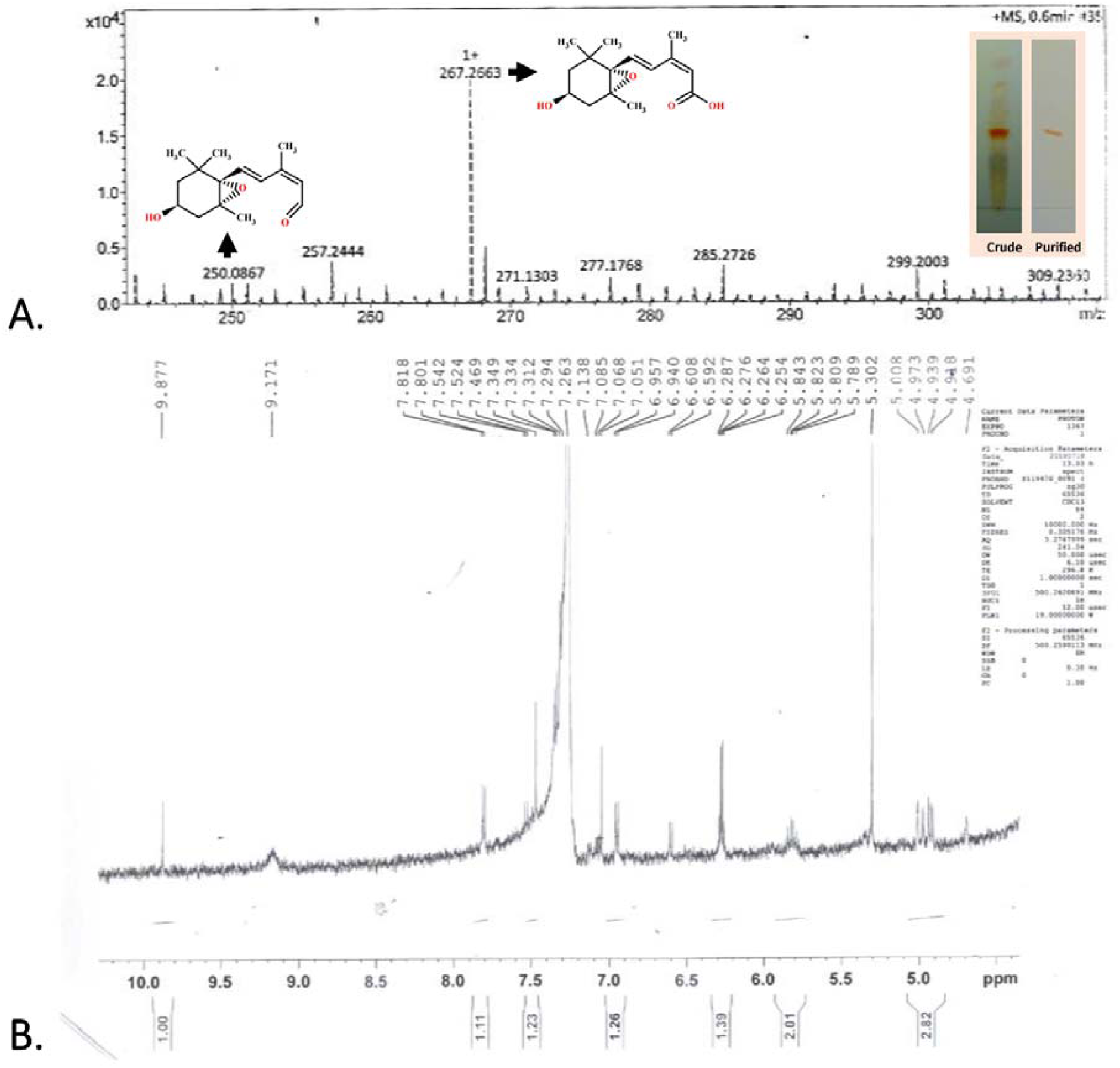
(A) HR-MS analysis of isolated F4 fraction (Insert figure represents crude and partially purified F4 from culture filtrate of CRDT-EB-1) and (B) ^1^H-NMR analysis of partially purified F4.

## 4. Discussion

The ABA is the prime hormone produced by the plants under abiotic stress, which regulates several physiological functions leading to the induction of drought stress tolerance in plants. Similarly, beneficial rhizobacteria also participate in the process of inducing abiotic stress tolerance by producing ACC deaminase through which ACC (the precursor of ethylene) is degraded, thereby reducing the ethylene level in plants (Glick, 2014). Additionally, recent studies discussed the possible involvement of bacterial synthesized ABA in mitigating drought stress in host plants (Cohen et al., 2009; Cohen et al., 2015). Perrig et al. (2007) reported that *Azospirillum brasilense* Az39 produces ABA as high as 0.077 μg mL^− 1^ in chemically defined Nfb medium. Similarly, ABA production was reported in many soils and plant endophytic bacteria such as *Lysinibacillus fusiformis* Ps7, *Bacillus subtilis* Ps8, *Brevibacterium halotolerans* Ps9, *Achromobacter xylosoxidans* Ps27 and *Pseudomonas putida* Ps30 (Sgroy et al., 2009), *Azospirillum brasilense* Sp 245 (Cohen et al., 2008) and *A. lipoferum* USA59b (Cohen et al., 2009). However, a detailed mechanism related to ABA production in bacteria and their involvement in inducing abiotic stress in host plant is yet to be studied.

In the present study, we isolated and characterized a beneficial rhizobacterium, which was significantly suppressed the drought stress in mustard seedlings under laboratory conditions. Upon characterization, it was found negative for both ABA and ACC deaminase production. Further, the drooping assay with different fractions of the bacterial extract revealed the existence of drought stress inducer molecule in ethyl acetate fractions of culture supernatant. By analyzing the results, we hypothesized that the selected bacteria might be producing certain metabolites other than ABA, which helps the host plant to withstand drought stress. Similar synthetic ABA-analogue B2, 8′-acetylene ABA, Cyano-cyclopropyl-ABA, etc. are reported to ameliorate abiotic stress and have similar biological functions in wheat, citrus, barley, etc. (Arbona et al., 2006; Frackenpohl et al., 2018; Zhou et al., 2019). Synthetic compounds “phosphonamide pyrabactin analogues” were found selectively elicit ABA-related responses such as stomatal closure and seed germination in plants (Van Overtveldt et al., 2015). Though, several synthetic structural and functional ABA-analogues were reported as ABA agonists, the occurrence of natural ABA analogues and their effects on the host plant to be studied.

Through TLC analysis, it was found that the F4 which separated at R_f_ value of 0.35-0.40 was found behaving similar to the ABA in function when analyzed through bioassays (stomatal closure, inhibition of seed germination, induction of drought stress tolerance, GA_3_ induced α-amylase inhibition in barley seeds and reversal of fluridon induced drought stress). Interestingly, *B. marisflavi* was showed the response to an increase in the concentration of stress inducers (PEG 6000 and NaCl) by producing a higher concentration of F4 in culture filtrate. In case of ABA production, similar observations were reported by Cohen et al. (2008) in the strain *A. brasilense* Sp245 where the strain found to produce a higher concentration of ABA when the osmotic potential (ψπ) of the medium was lowered from −0.2 to −0.7 MPa by the addition of 100 mM NaCl. Furthermore, endophytic bacteria isolated from sunflower showed similar behavior by responding to increase concentration of PEG 6000 (lowering ψπ from 0 to –2.03 MPa) through increased production of ABA in culture media (Forchetti et al., 2007). In our study, the characterization of the purified compound revealed the presence of xanthoxin and xanthoxic acid, which are well-known precursors of ABA in plants. Among which xanthoxin is unstable, it may convert into a more stable xanthoxic acid form. However, the involvement of some of the minor compounds in F4 which were unidentified, in inducing resistance against drought stress cannot be ignored.

The results supported our hypothesis strongly suggesting the involvement of bacterial secreted compounds other than ABA in suppressing abiotic stress in the host plant. In contrast to five stimulatory plant hormones, xanthoxin is an intermediate in the biosynthesis of ABA in plants (Seo and Koshiba, 2002) and known as an endogenous plant growth inhibitor (Firn and Friend, 1972; Taylor and Burden, 1970a; Taylor and Burden, 1970b). Interesting facts are that the absolute structural configuration of xanthoxin is very much similar to ABA and it plays a similar role of ABA in its physiological activity (Burden et al., 1971; Taylor and Burden, 1970b). Xanthoxin is widely occurs in various plant species (Firn et al., 1971) and responsible for stomatal closure (Raschke et al., 1975). It can be formed by the photo-oxidation of violaxanthin and also acts as a seed germination inhibitor (Burden and Taylor, 1970; Taylor, 1968; Taylor and Burden, 1970b; Taylor and Burden, 1972; Taylor and Smith, 1967). The similarity in xanthoxin and ABA function was established by Taylor and Burden (1970a), where they identified two natural plant growth inhibitors from dwarf-bean and wheat, and identified them as two isomers derived through photo-oxidation of xanthophyll, i.e., xanthoxin. Through bioassay, they observed that the function of one of the isomers *cis*- and *trans*-configurations of xanthoxin is similar to that of ABA in plants. Also, an increased concentration of ABA in tomato seedlings treated with xanthoxin (Taylor and Burden, 1972).

In plants, ABA biosynthesis begins normally inside the chloroplast, starting with oxidative cleavage of carotenoids, 9-*cis*-violaxanthin or 9-*cis*-neoxanthin to xanthoxin by the plastid enzymes 9-*cis*-epoxycarotenoid dioxygenases (NCEDs). Xanthoxin is secreted into the cytoplasm, where it is further oxidized with the help of xanthoxin oxidase to abscisic aldehyde. In the final step, the abscisic aldehyde is converted to ABA by the action of abscisic aldehyde oxidase (reviewed by Taylor et al., 2000). During the course of evolution, as per the endo-symbiotic theory, the photosynthetic bacteria are believed to be engulfed by non-photosynthetic eukaryotes, thereby attaining the ability of photosynthesis. Further, this association is continued to evolve, which made the existence of all plant species possible on earth. As a relic of this evolutionary past, the chloroplast still contains its genomes and protein synthesis machinery.

No ABA biosynthesis pathway is currently described in bacteria. Considering the above-mentioned information, it was hypothesized that *B. marisflavi* might follow either the same or different pathway of chloroplast to synthesize ABA-analogue or xanthoxin. It is, however, possible that different prokaryotic species utilize independently evolved carotenoid-dependent and -independent ABA biosynthesis pathways (Lievens et al., 2017). Also, *Achromobacter, Bacillus* and *Pseudomonas* are all known to produce carotenoids, which make them likely that these bacteria follow a carotenoid-dependent pathway for the production of ABA-analogue. *Bacillus marisflavi* is well studied for its carotenoid biosynthesis (Khaneja et al., 2010). Further, in our study, it was observed that when CRDT-EB-1 was exposed to drought stress, it produced F4 containing xanthoxin at a significantly higher level. These observations open a new avenue to study xanthoxin biosynthesis pathway in *B. marisflavi*, which may be different from plant pathways.

## 5. Conclusions

Here, it was hypothesized that *B. marisflavi* at rhizosphere under drought stress conditions might catabolize the carotenoid to produce ABA analogue/ xanthoxin. This low molecular compound may be up taken by plants where it may stay in its original form or further converts into ABA with the help of xanthoxin oxidase and abscisic aldehyde oxidase. Further, they induce the plant to undergo physiological adaptation against drought stress. As per our knowledge and literature survey, the outcome of this study is the first report of natural bacterial synthesized ABA analogue/ xanthoxin and their involvement in imparting resistance against drought stress in host plant.

## Author contributions

GHG carried out laboratory experiments including morphological, physiological and biochemical parameters of the seeds/ plants subjected to various treatments. DP and AS characterized bacteria for their beneficial traits and identified. SD performed the HPLC, LC-MS, HR-MS and NMR studies. SLG compiled the chemistry data and identified possible bioactive compounds. HP designed the research and experimental work. SRN and HP were written the manuscript. All authors are equally contributed to writing paper and, read and approved the final manuscript.

## Acknowledgements

The authors are thankful to Indian Institute of Technology Delhi, New Delhi, and University of Mysore, Mysuru for facilitating this investigation.

## Supplementary data

**Fig. S1.** (A) Morphology on nutrient agar and (B) A phylogenic analysis of 16S rRNA gene of *Bacillus marisflavi* CRDT-EB-1.

**Fig. S2.** (A) Thin layer chromatography (TLC) analysis and (B) High performance liquid chromatography (HPLC) of crude extract of culture filtrate of CRDT-EB-1 for ABA detection.

**Fig. S3.** Liquid chromatography-Mass spectroscopic analysis of (A) culture filtrate of CRDT-EB-1 and (B) ABA standard.

**Fig. S4.** Effect of abiotic stress on extracellular metabolites production by CRDT-EB-1. **Table S1** Morphological, physiological and biochemical test results of *B. marisflavi* CRDT-EB-1.

