## Supplementary files for "ABA analogue produced by *Bacillus marisflavi* modulates the physiological response of host-plant under drought stress"

**Supplementary data**

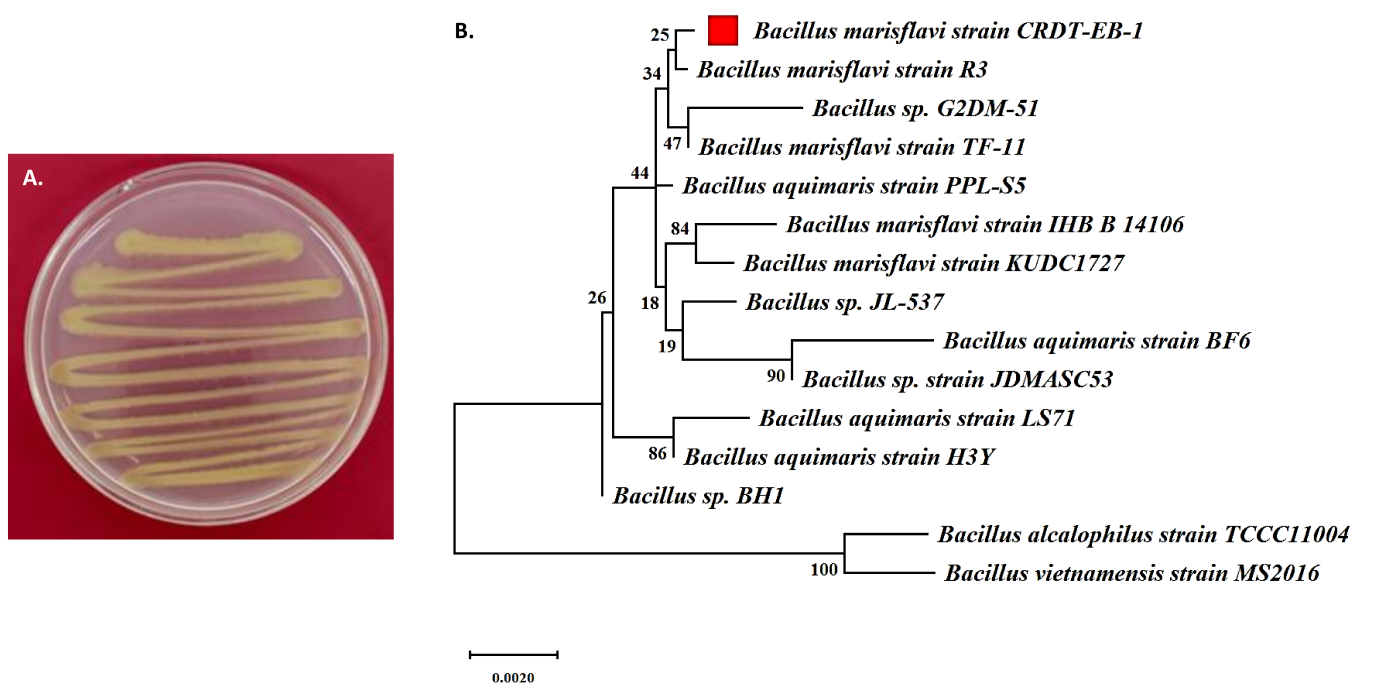

**Fig. S1.** (A) Morphology on nutrient agar and (B) A phylogenic analysis of 16S rRNA gene of *Bacillus marisflavi* CRDT-EB-1.

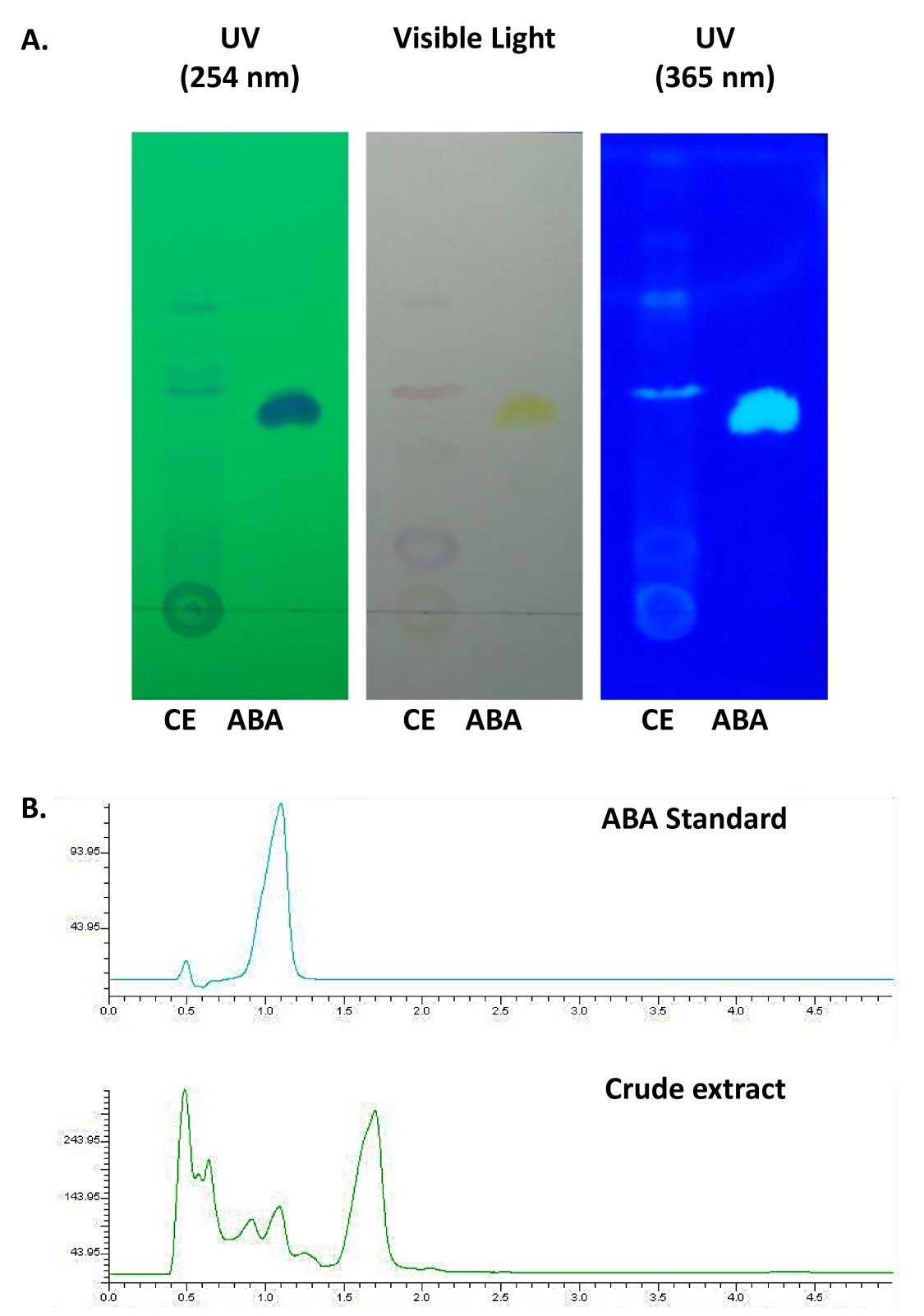

**Fig. S2.** (A) Thin layer chromatography (TLC) analysis and (B) High performance liquid chromatography (HPLC) of crude extract of culture filtrate of CRDT-EB-1 for ABA detection.

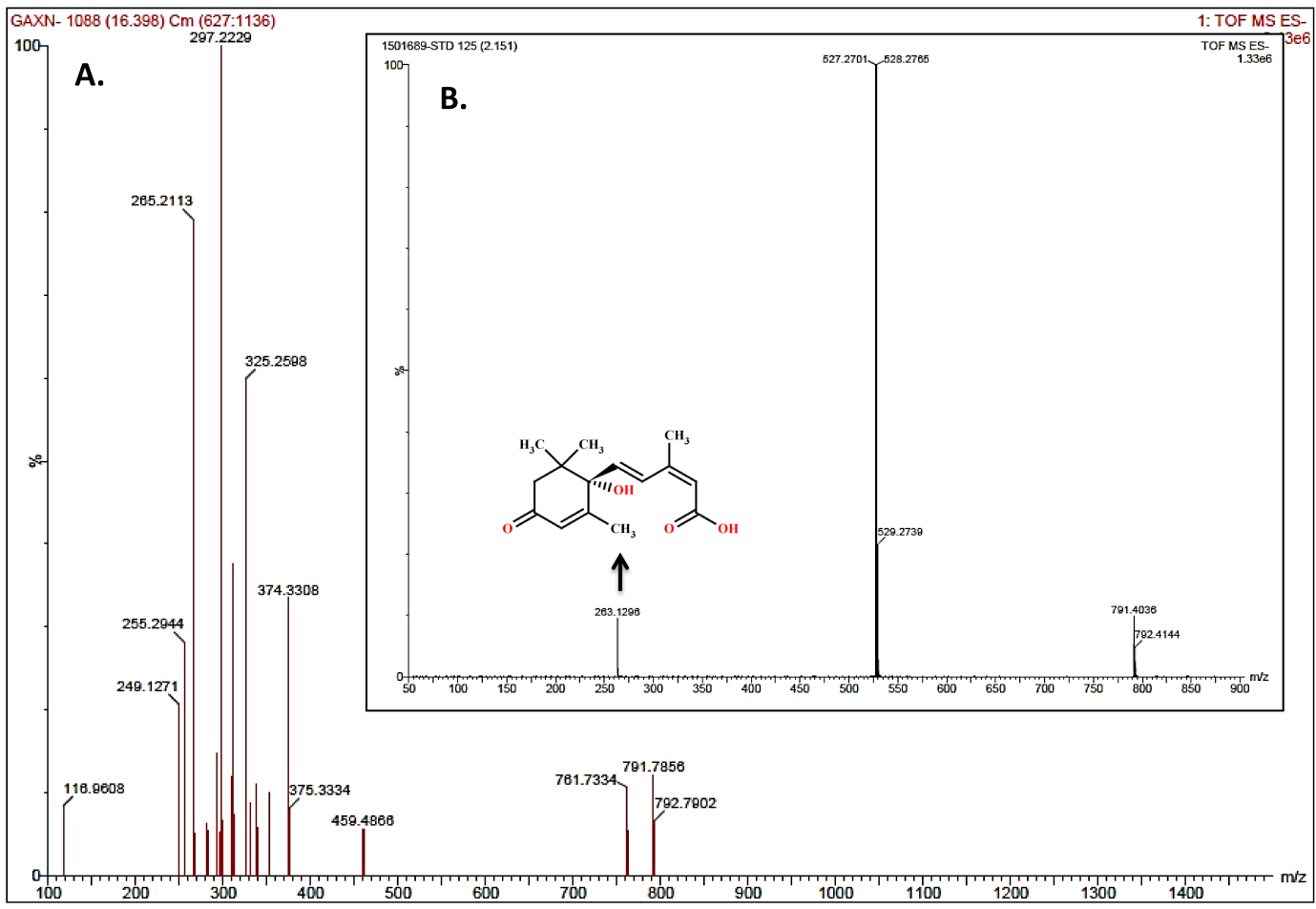

**Fig. S3.** Liquid chromatography-Mass spectroscopic analysis of (A) culture filtrate of CRDT-EB-1 and (B) ABA standard.

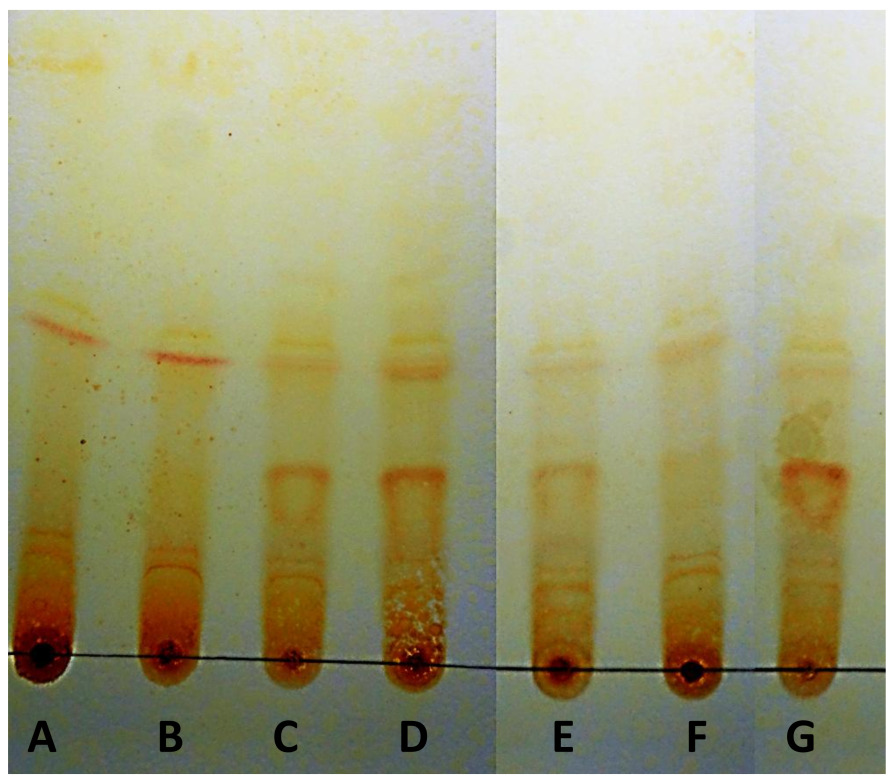

**Fig. S4.** Effect of abiotic stress on extracellular metabolites production by CRDT-EB-1. (A) defined media, (B) defined media amended with 2.5% PEG 6000 and 2.5% NaCl, (C) defined media amended with 5% PEG 6000 and 5% NaCl, (D) defined media amended with 7.5% PEG 6000 and 7.5% NaCl, (E) defined media amended with 10% PEG 6000, (F) defined media amended with 8% NaCl and (G) defined media amended with 10% PEG 6000 and 8% NaCl. All the defined media also amended with 5 g L^–1^ beef extract, 5 g L^–1^ glucose and 2.5 g L^–1^ peptone. The TLC plates were visualized after spraying with 2,4-dinitrophenylhydrazine reagent.

**Table S1** Morphological, physiological and biochemical test results of *B. marisflavi* CRDT-EB-1.

| **Sl. No.** | **Character/ Test** | **Result** |
| --- | --- | --- |
| 1. | Gram staining | Gram +ve (Gram-Variable) |
| 2. | Spore position | Central/ Sub-terminal endospores |
| 3. | Colony color | Dark to pale yellow |
| 4. | Oxygen requirement | Aerobic |
| 5. | Growth range (°C) | 10-45 |
| 6. | Maximum growth at (°C) | 30-37 |
| 7. | Growth range (pH) | 3-11 |
| 8. | Maximum growth at (pH) | 5-9 |
| 9. | Growth range (% NaCl) | 0-15 |
| 10. | Maximum growth at (% NaCl) | 0-8 |
| 11. | Growth range (% PEG 6000) | 0-25 |
| 12. | Maximum growth at (% PEG 6000) | 0-10 |
| 13. | ONPG | – |
| 14. | Lysine utilization | + |
| 15. | Ornithine utilization | + |
| 16. | Urease | – |
| 17. | Phenylalanine Deamination | V |
| 18. | Nitrate reduction | V |
| 19. | H_2_S production | – |
| 20. | Citrate utilization | – |
| 21. | Voges Proskauer's | – |
| 22. | Methyl red | – |
| 23. | Indole | – |
| 24. | Malonate utilization | – |
| 25. | Esculin hydrolysis | + |
| 26. | Arabinose | – |
| 27. | Xylose | – |
| 28. | Adonitol | – |
| 29. | Rhamnose | – |
| 30. | Cellobiose | + |
| 31. | Melibiose | + |
| 32. | Saccharose | + |
| 33. | Raffinose | + |
| 34. | Trehalose | + |
| 35. | Glucose | + |
| 36. | Lactose | – |
| 37. | Oxidase | – |

Note: ‘+’ indicates positive; ‘–’ indicates negative.
